# Current challenges in microbiome metadata collection

**DOI:** 10.1101/2021.05.05.442781

**Authors:** Gabriel Rosenfeld, Angelina Angelova, Chris Shin, Mariam Quinones, Darrell Hurt

## Abstract

While the biomedical community has embraced data sharing (e.g. results, raw data) and supported establishment of large research consortia (e.g. the Human Microbiome Project) aimed to standardize the quality of important sets of microbiome sequencing data, the reusability of most microbiome data is still limited by the quality of its associated metadata. To ensure that microbiome data is indeed FAIR (Findable, Accessible, Interoperable, and Reusable), it is necessary to consider tools and approaches that make it easier to provide high-quality metadata that is fit for purpose moving forward. Such tools and approaches could be informed by current efforts to harmonize and improve the quality of extant microbiome metadata.

## Introduction

The scale of microbiome data being shared by the community is rapidly growing. An NCBI BioSample search for “((metagenome OR microbiome)) AND human” resulted in over 600,000 records in January 2021. These numbers will only increase over time and providers of public microbiome data (e.g. EBI, DDBJ, NCBI) are simultaneously developing useful interfaces, APIs, standardized analytical pipelines, and other tools to make it easier to search and access shared datasets. As of December 2020, the MGnify database which provides a set of such APIs and standardized pipelines, contained over 140,000 samples just from human hosts (1). The Genomics Standards Consortium and other community initiatives work in close collaboration with providers of public microbiome data, to ensure community-wide implementation of data standards and facilitate the sharing and reuse of data (2). Nevertheless, the reusability of this shared scientific data is impeded by the current state of the accompanying metadata.

In this article, we demonstrate the challenges we encountered when trying to use sample metadata and the barriers it creates for data reuse as an example case study. Our goal was to leverage public microbiome profiling data to develop a way to compare the relative increase or decrease in microbial taxa by way of metadata comparators (e.g. diet, age, gender, ethnicity) across human studies. Unfortunately, our efforts were quickly hindered due to the encountered challenges of trying to harmonize and rectify missingness of sample metadata. As a way for others to directly assess our code, findings, and data, we include a link to a Figshare repository (https://doi.org/10.6084/m9.figshare.13918190.v1). We also include an interactive html report (human_sample_metadata_missingness.html) within this Figshare repository under the reports folder, which is referenced extensively below. We chose this format to be transparent and allow others to explore the challenges of metadata curation and specific problems of interest to them. The html file can be opened in most modern web browsers and the filtered missingness table can be downloaded and exported in a variety of formats.

### Observations of Metadata from Putative Human Samples

Table #1 in the interactive html report demonstrates available metadata for samples (presumed to be of human host based on record matches of column names indicative of host organism). Table #1 shows over 3000 different column descriptors illustrating the wide use in terminology used to describe common elements of sample metadata across publicly shared human studies. While some columns could reflect additional data elements from non-human samples that were included in the dataset, the majority of data types of interest (e.g. age, gender, ethnicity) exhibit multiple columns containing similar data types. The abundance of terms used to describe similar variables presents challenges for data-driven approaches by introducing a need for extensive manual curation efforts to harmonize across metadata. Such variability is not isolated due to the specific nature or design of each microbiome study (e.g. environmental, disease, diet, etc.). Rather, it appears to reflect systematic issues resulting from current data submission practice where researchers have to map individual metadata fields of the study to community standards while balancing modern demands of biomedical research such as publication requirements and privacy issues.

Aside from the organizational inconsistencies within the metadata, Table #1 also demonstrates a significant level of sparsity (missingness/absence) of basic information associated with these human microbiome studies (e.g. host species, host gender, sample type [control/experimental], sampling site [e.g. skin/gut], etc.). Table #1 shows that certain attributes matching the pattern “age” contain more than 81.6 % missing values (e.g. “characteristic.host.age”, “characteristics.host.age.unit” and other variations of the “age” descriptor). Exploring the location of host sample site (basic information needed to compare across samples or studies), a similar trend of high missingness is observed. The characteristic “host_tissue_sampled” presents with 94% missing values, while characteristics such as “body.habitat”, body.product”, and “body.site” attempting to describe a similar data type produced missingness of ∼82% for each characteristic. This lack of consensus for descriptive terminology in microbiome studies and high-level of missingness create barriers to metadata mining and impedes meta-analysis approaches. For instance, “host.age”, “host.sampling.site”, or “host.gender” appear as relatively straightforward column descriptors that could describe a human microbiome dataset yet are rarely populated based upon the observations above or are deposited in a variety of metadata columns using non-standardized terms. The associated Figures 1-4 show visualizations of missingness for sets of metadata columns related to age, body mass index, ethnicity, and gender, which were types of metadata of interest for our original analytic goals.

**Figure 1.**
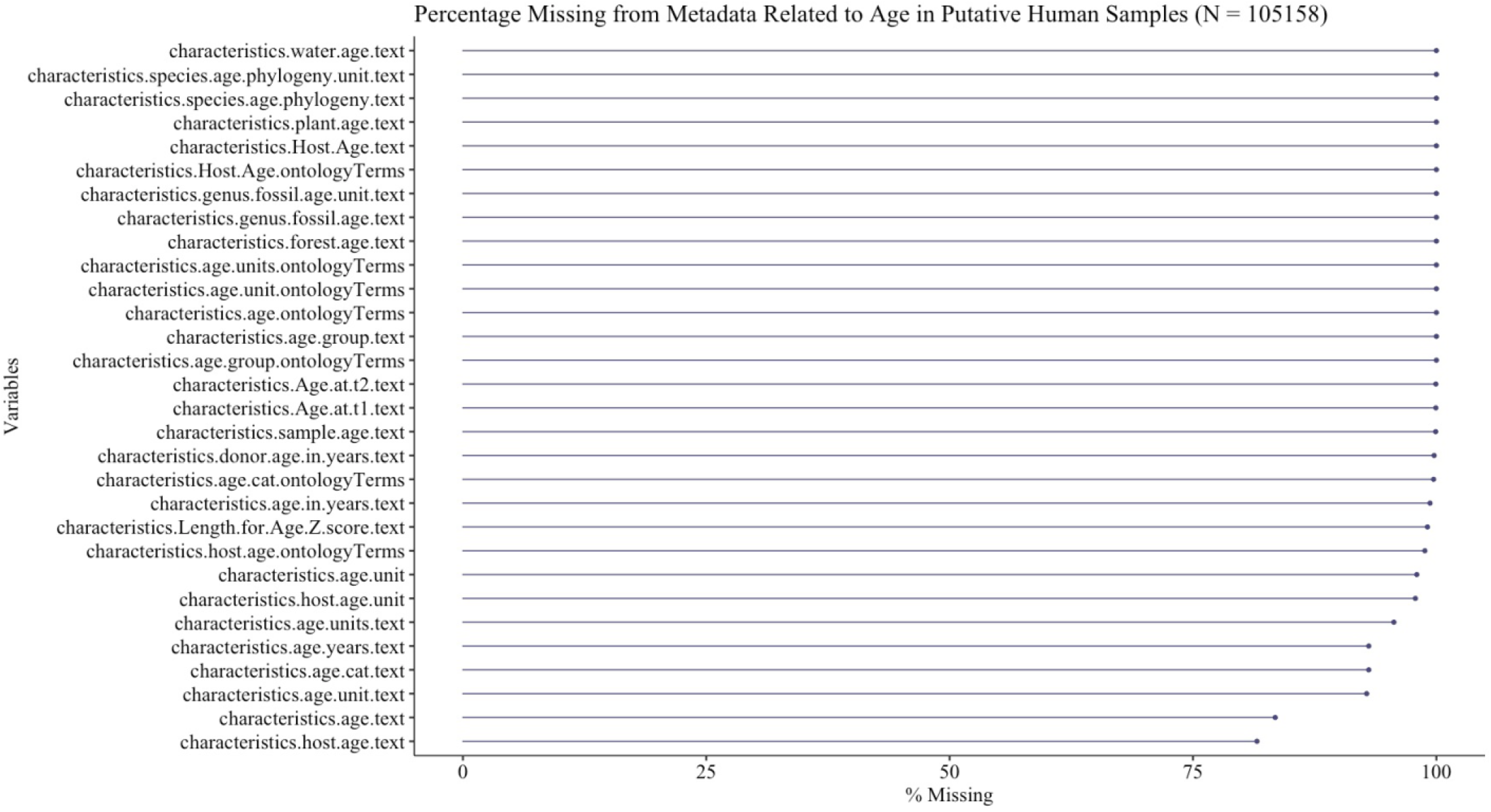
Metadata column missingness is shown for columns relating to age sorted by percentage missingness. The 3000+ metadata columns associated with putative human amplicon microbiome samples were filtered for those matching an age-related regex expression for visualization using the “naniar” R package.

**Figure 2.**
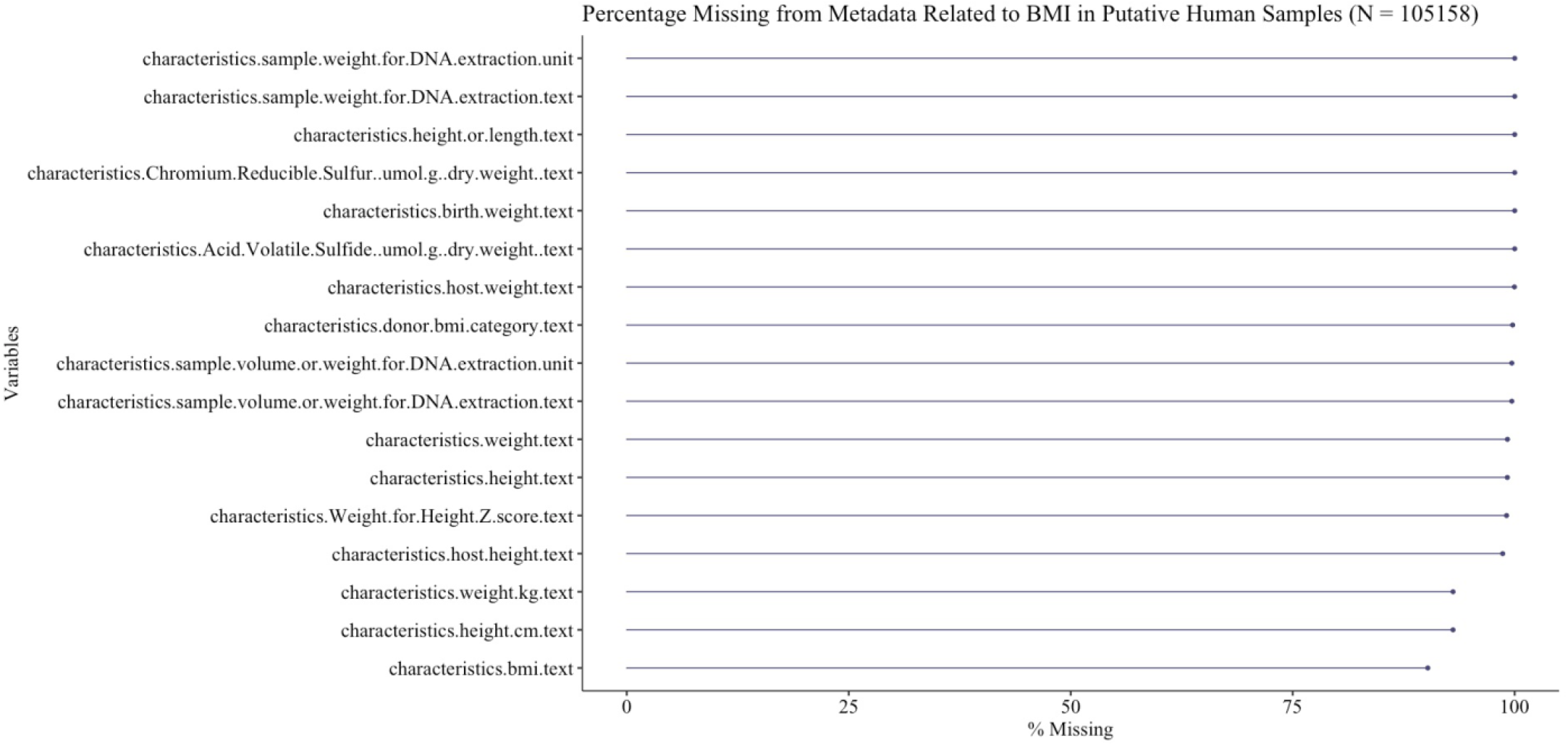
Metadata column missingness is shown for columns relating to body mass index sorted by percentage missingness. The 3000+ metadata columns associated with putative human amplicon microbiome samples were filtered for those matching an BMI-related regex expression for visualization using the “naniar” R package.

**Figure 3.**
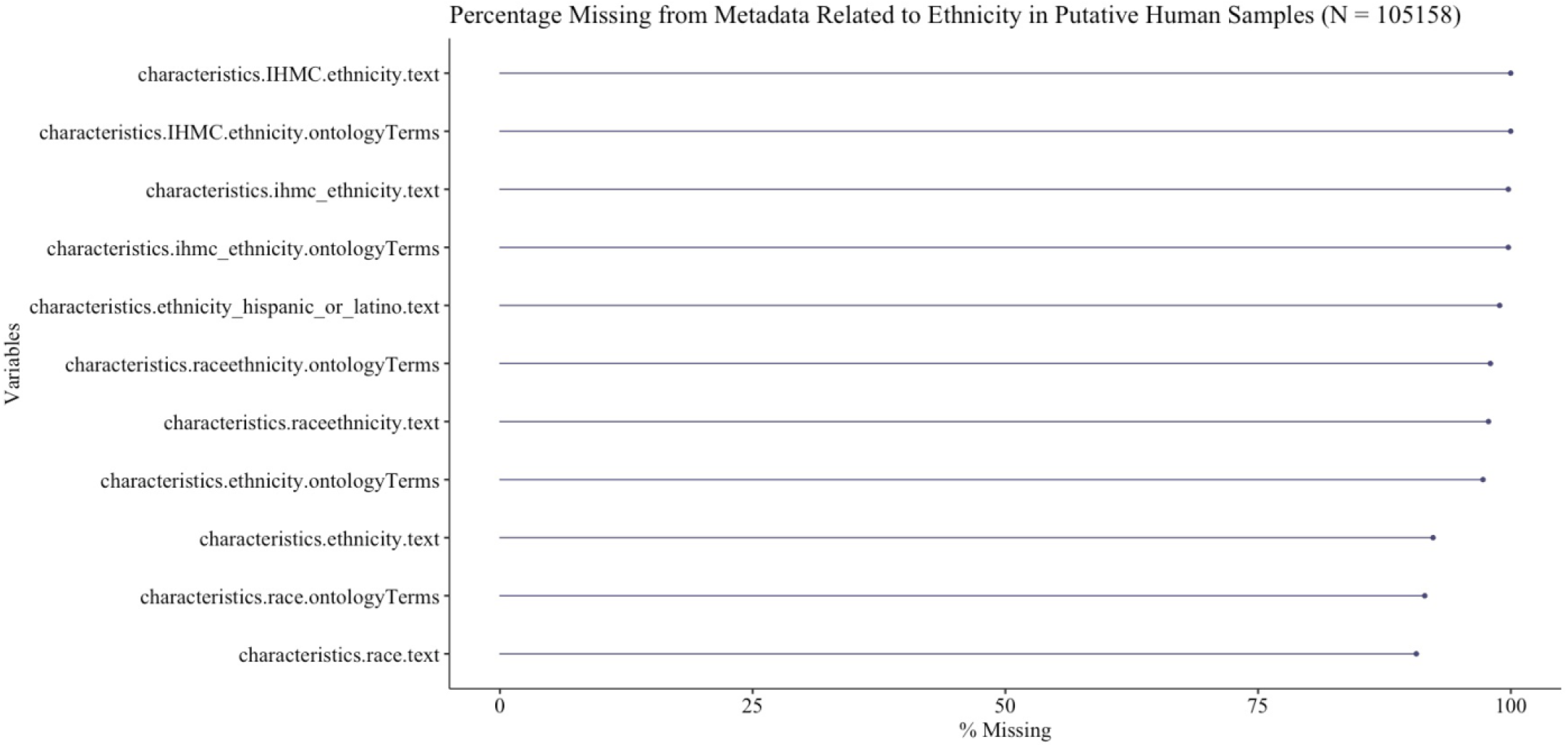
Metadata column missingness is shown for columns relating to ethnicity sorted by percentage missingness. The 3000+ metadata columns associated with putative human amplicon microbiome samples were filtered for those matching an ethnicity-related regex expression for visualization using the “naniar” R package.

**Figure 4.**
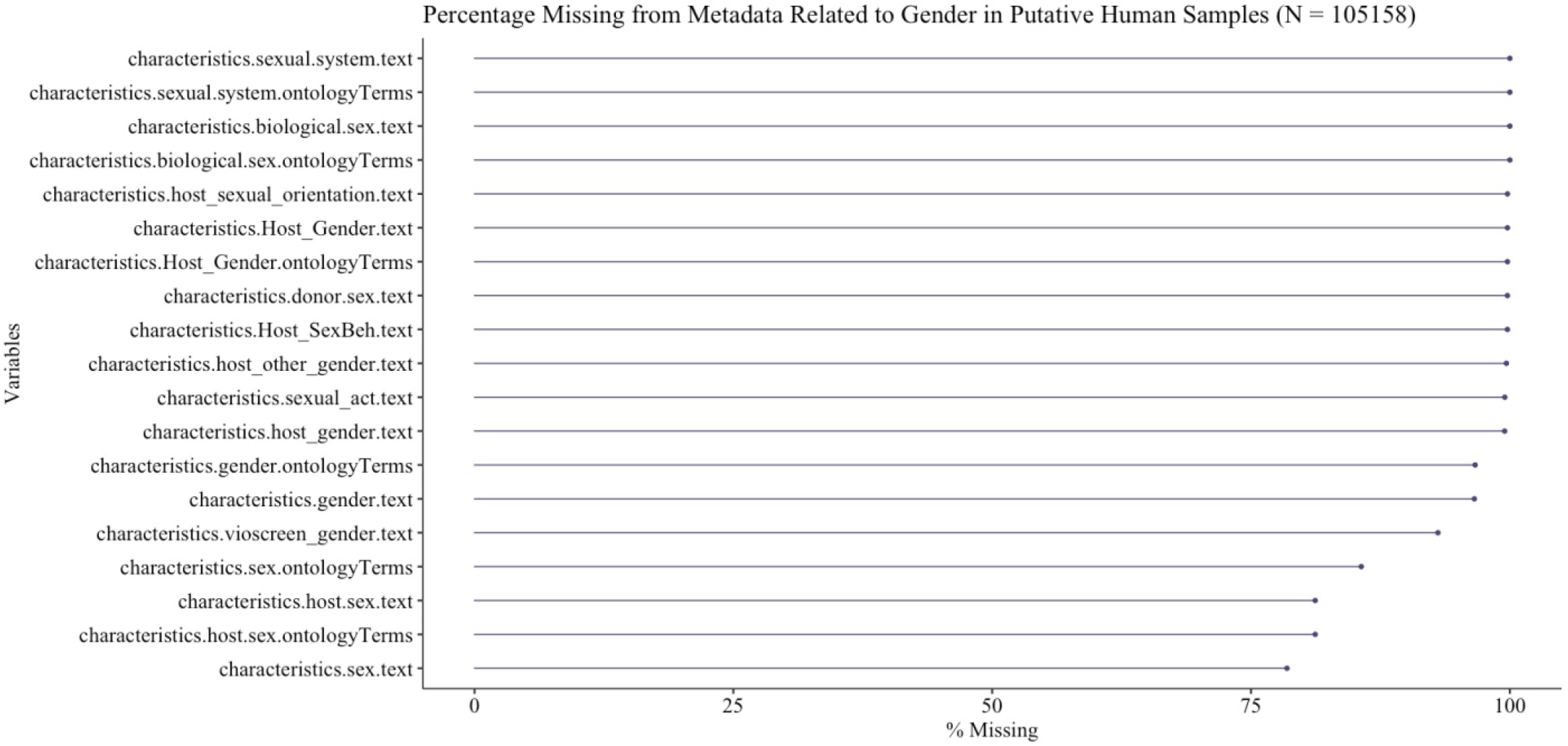
Metadata column missingness is shown for columns relating to gender sorted by percentage missingness. The 3000+ metadata columns associated with putative human amplicon microbiome samples were filtered for those matching a gender-related regex expression for visualization using the “naniar” R package.

Sparsity or missingness is lower if metadata elements relating to a feature of interest (e.g. age, gender, sample site) are harmonized across studies; nonetheless, such requirements slow the pace of data-driven research. Moreover, outside of obvious descriptors such as age and gender, it become difficult to ascertain exactly which columns should be combined and necessitates sometimes highly complex filters to select the right set of columns. For example, in the interactive report, we demonstrate the challenge of harmonizing the attribute “age”, which seemed a relatively straightforward task. In Table #2 of the html report, you can see the level of missingness in intermediary and harmonized age metadata columns. Concerningly, the “any_age” variable corresponding to records with any age related column information (identified using regex matching), showed only ∼40% completeness. When trying to harmonize using available metadata, only ∼18% of records could be harmonized to a common age (age_derived variable). This is because either the age or age_units information was missing or was non-standardized making it difficult to harmonize.

When available, the community-wide selection and application of consensus descriptors or common data elements (CDEs) to describe a study can significantly reduce the activation energy needed to repurpose the data. CDEs can facilitate the development of automated approaches for harmonizing the metadata and increasing its utility (3) and the National Library of Medicine offers a service for searching CDEs (https://cde.nlm.nih.gov/cde/search). The issue of lack of consensus descriptors becomes even more challenging when considering information that would be specific for each study. We were curious about whether samples came from smokers or non-smokers and investigated the columns where this information might be found. In one study, the characteristic, “environment.feature”, was found to contain information about the host’s smoking habits. Uncertainty about where the data should be provided contributes to lack of metadata compatibility, presents challenges for data cleanup or rescue, and reduces power from secondary studies interested in samples with these attributes. A worst case scenario would involve incorrect comparisons being made due to confusion about the metadata. In the case of host smoking status being shared in the “environment.feature” column, it suggests confusion with mapping to the community-standard to use for describing smoking. According to the mixs_v5 template provided by GSC, it would likely be more appropriate to use the “smoker” column (http://press3.mcs.anl.gov/gensc/files/2020/02/mixs_v5.xlsx). We hope that these examples demonstrate some of the complexity of the issues created by the current state of metadata which preclude maximal reuse of publicly deposited metagenomic data.

### The Systematic Issues Involving Data Sharing

Our observations of the challenges we encountered with metadata quality suggest a systematic issue in current data sharing practices. These suggestions are not intended to cast blame at any one of the individual contributors to the data sharing process such as the data depositor or repositories. We appreciate the pressures of publishing and the additional demands that depositing data places on all involved. Rather, they are presented to raise awareness of the subsequent lack of metadata quality resulting from the current practices in place. And hopefully by highlighting the issue, we can be motivated as a community to identify potential areas where the process could be improved. Alas, the admirable goals of seeing that this data could be maximally reused by others including the original data depositor do not seem feasible if current practices remain. An important part of this process will be a way to build trust amongst all interested parties; for example, those interested in data reuse (including the authors here) have to be more vocal in how they anticipate reusing the data to ascertain exactly what should be captured to maximize reusability and minimize burden on the data submitter.

The observed issues in metadata quality and completeness, could be categorized in a few groups: I) analogous information being split over multiple fields of the metadata II) the information is recorded using non-standardized/variable vocabulary/terminology, III) the information does not vary within the study and therefore is not recorded/provided in the metadata (e.g. host organism used), IV) information for specific samples is omitted, V) the information does not fit conventional/common data descriptors and is therefore omitted or provided/coerced into non-conventional descriptors (rather than the specific descriptors being added or edited).

Some of the systemic reasons leading to the above issues reflect the realities of modern biomedical research including the pressure to publish, the transience of the biomedical workforce where a project might be started by one researcher and completed by another, the privacy aspects of health-related data, and a potential misunderstanding of how the data might be repurposed by secondary data users. This leads to current practices such as sharing only minimally required data despite additional metadata being potentially useful or even necessary for a complete picture of how the data were collected and used within the study. Due to these thorny issues, researchers rarely provide information beyond the minimal deemed required for study reproducibility. For example, if researchers have not explored the effect of a covariate or have kept such a factor consistent across the samples of their metagenomic study (collect samples from only male subjects for example), they may not include such descriptors in their metadata despite its fundamental nature. Unfortunately, such practice diminishes the utility of the deposited data for repurposing (e.g. a meta-analysis considering the effect of gender on microbiome communities) by contributing to the problem of informational “missingness” within and between datasets. More insidiously, competitive pressure can create an incentive not to fully describe a dataset if samples may be used for future analyses or publications. Whether warranted or not, data submitters may fear being scooped. Secondary studies are often considering approaches that would leverage the already published hypothesis and results such as a meta-analysis or similar methodology to increase statistical power of detecting effects. Given that those interested in reusing data are not aware of the hypothesis, goals and analytical methods in the forthcoming studies, it may be worth a community-wide discussion of the potential risk of data sharing against the cost toward repurposing the data for new science.

### Further discussion and potential opportunities

Our experience with the microbiome metadata harmonization problem, lead us to consider the reasoning behind collecting such data in the first place by the scientific community. What should determine a metadata’s fitness for purpose in other words. The initial idea behind collecting such data began with the necessity to make it possible for a study to be reproduced to confirm its results (and metadata should be meeting this standard at a minimum). With increasing size of publicly available datasets, however, the scientific community begun to shift its interest away from reproducing the analysis of another study and more towards comparative analyses of their dataset against others as well as combining studies through meta-analyses. Initiatives such as the Genomic Standards Consortium (GSC) formed as community efforts to facilitate these new requirements of publicly-shared genomic datasets (2). Recently, the computational infrastructure has advanced (e.g. compute clusters, cloud providers, GPUs, cheap data storage) to support ever larger cross-study analyses involving genomic data including metagenomics as part of “big data” initiatives.

Enforcement of standards or quality at the time of submission has been limited historically, which may be indicative of the challenges involved in their enforcement (cost, time, etc.) as well as the cost/benefit ratio given recent computational advances discussed earlier. The new NIH Final Data Management Policy (https://grants.nih.gov/grants/guide/notice-files/NOT-OD-21-013.html) makes it clear that all data and associated information (e.g. the metadata) needs to be of sufficient quality to support reproducibility of the study; therefore, it remains to be seen how such policies may impact submission moving forward. More importantly, enforcement is only one tool in the toolbox and likely will not work well in the absence of education and spreading awareness amongst the scientific community. Promulgating best practices for data sharing as well as the ways in which publicly shared data is likely to be used can be an effective part of a strategy to improve future metadata quality. By highlighting that public data is unlikely to be used in a way that would jeopardize data submitters (e.g. being scoped) and more likely to be used to expand scientific inquiry that may directly benefit the original data depositor, trust can be built among data providers, and those interested in reusing the data.

As part of the trust-building, sufficient detail and standardization of submitted metadata to meet minimum study-level reproducibility should be the goal of data providers moving forward; however, fundamental study-wide descriptive information for each sample that could elucidate the study design should be shared as well whenever feasible. In terms of host-microbiome data, this may include (but is not limited to) host species, site of sampling from the host (e.g. skin, lumen), type of microbiome analysis (e.g. amplicon, metagenome, meta-transcriptome), targeted sequence (e.g. V3-V4 rRNA region), etc., even if such factors are consistent throughout the samples. Many of these types of information are already recommended and are customized for the different types of studies as suggested by the Genomics Standard Consortia. In short, the community is continuously assessing, updating, and recommending appropriate metadata that should be included for each type of study (e.g. http://www.obofoundry.org/, https://gensc.org/) Nevertheless, in practice, these standards are still not always being adhered to. While greater trust can be built by data submitters with greater compliance to best practices, the responsibility cannot just be placed on them. Those interested in reusing the data will have to engage with submitters and repositories to demonstrate how they plan to reuse the data. And it will be necessary to identify ways to make the data submission process easier including how to share metadata that is of sufficient quality and standards to maximize data reuse.

As an example, submission guiding tools are being developed to aid the submitter through the appropriate metadata organizational and descriptive strategy during the submission process. Tools such as METAGENOTE (4) (https://metagenote.niaid.nih.gov/) aim to facilitate data submission to correctly classify and organize data and associated metadata by recommending standardized columns for metadata associated with experimental design or focus. For instance, in human host gut longitudinal studies with interventions, columns such as ethnicity of host, comorbidities, diet or time points for sample collection are suggested. On the other hand, in human host lung observational non-longitudinal experimental designs, different columns are recommended more applicable to that type of experiment (e.g. host smoking status, observation group, age, etc.).

Development of machine learning (ML)-based tools to guide data submission are another strategy that could facilitate improvements in metadata quality. One example would be data ingestion assistance agents that make real-time recommendations for needed information as well as steps that check data deposition and prompt data depositors on potential discrepancies in the data. Such tools might not only guide data submitters but can also learn from them the kind of additional metadata that researchers might be providing in order to describe their data. With such knowledge, the tools can improve their suggestions to future submitters. ML tools might warn submitters against incorrect, incomplete data descriptors to support improvement in the correct use of metadata column headers, but also the metadata entries themselves, reducing the barrier to (meta)data reuse by decreasing subsequent curation or harmonization efforts. The Center for Expanded Data Annotation and Retrieval (CEDAR) serves as a an example of the type of model that could be applied in real-time to metadata (5-7) and tools such as BioPortal demonstrate how metadata could be mapped to common ontologies (8). As the scientific community gains experience with the requirements for the minimum viable metadata for the reusability and reproduction of the studies, it will hopefully become easier on the data depositor to select standardized vocabularies and ontologies for effective data description.

Our experience with exploring large-scale metadata sets have convinced us that it is time for our scientific community to formalize the minimal required metadata necessary to reproduce at least the main findings of the original study. The increasing efforts in cross-study analyses (meta-analysis) efforts would allow us to familiarize ourselves with the type of minimal metadata required for larger-scale analyses. Optimally, the goal would be for an external data accessor to obtain a basic understanding of the design and sample source and organization of each study, without having to invest significant time reading the associated publication. We believe metadata could accurately reflect the high-level study design and data-driven efforts would have to rely on a (meta)data-centric approach to be scalable. Standardizing metadata submission procedures for this purpose would require more stringent informational demands on metadata per sample (e.g. columns for type of experimental design, experimental group, time frame [with units], microbiome data type, etc.) to allow for more diverse utilization of the submitted datasets. The GSC provides templates that likely already cover such information. Data quality mindfulness is increasingly vital and with it the necessity for tutorials, articles and protocols that could guide the data submitters through the correct submission process along with the submission tools.

Community driven efforts both internal and external to NIH can be informative when considering strategies or approaches to improve future metadata. Several community driven resources such as MG-RAST (9, 10) and Qiita (11) have been collecting microbiome data and metadata in recent years. Examples of records from each database demonstrate the breadth of metadata being collected across sample sites such as skin, gut, and lung (see Metadata Examples.xslx in the Figshare supporting files). While this paper indicates there are impediments to the automated use of microbiome data due to metadata, research groups are putting focus into collecting and utilizing data that meets sufficient quality and interest. Though there are challenges, some progress is being made with its reorganization, harmonization and successful exploration. For example, the IHD database leverages stool microbiome data across independent studies with cases and controls for a meta-analysis (2). The authors were then able to also examine the effects of various batch normalization tools pooling samples across diseases and increasing the statistical power of differential OTU detection (3). Also, the Integrative Human Microbiome Project has undertaken a consortia-wide effort to collect the required metadata to support data reuse and repurposing (12). Such projects serve as an example of the type of work that could be better supported with quality improvements in metadata. The National Microbiome Collaborative is working with the community to specifically address data reuse including metadata quality (13, 14).

The Bioinformatics and Computational Biosciences Branch within NIAID works on the development of community-driven tools to assist in the generation and management of microbiome data such as METAGENOTE (4) (https://metagenote.niaid.nih.gov/) and Nephele (15) (https://nephele.niaid.nih.gov/index) and is interested in helping facilitate these community efforts. We believe the message of this article is timely given the recent release of the Final NIH Policy for Data Management and Sharing (https://grants.nih.gov/grants/guide/notice-files/NOT-OD-21-013.html), which encourages the sharing of data to support reproducibility and transparency of NIH-supported research. The demand for reproducibility is not just important from the aspect of the individual study itself but also helps organize the required metadata for data repurposing to support new science.

Despite community efforts to generate and adhere to standards, implementing such standards is not straightforward process. Due to the variety of experimental types and possible designs, the community is already immersed with both usable and unusable data. It is therefore crucial to start investing time and effort into exploring solutions to make data re-utilization easier and more efficient for the entire community. One of the most productive approaches will be to take a hard look at the initial submission step where (meta)data quality issues first arise. It will take a team effort involving data generators, secondary data users, data repositories, and other interested stakeholders in the data ecosystem to build trust and understanding on how best to encourage maximal data reuse with minimal level of burden.

## Materials and Methods

### Computational Environment

This work utilized the computational resources of the NIH HPC Biowulf cluster. (http://hpc.nih.gov).

### Code and data used for the analysis

#### Code

The analysis was completed as a drake workflow in R. The code for the analysis demonstrating all steps of the pipeline can be obtained as part of the supplementary data in Figshare (https://doi.org/10.6084/m9.figshare.13918190.v1).

#### Data

Data were pulled directly from Mgnify or Biosamples resources using available APIs. Data were pulled from each API in summer 2020; therefore, the individual files in the downloads folder provide a time stamp of when the data were last pulled. The API calls focused on using analyses from MGnify pipeline version 4.1 for amplicon analyses. Relevant metadata files can be found in the data and downloads folders in the unzipped supplementary file found in the Figshare link.

#### Interactive HTML report

The interactive HTML report is provided as part of the supplementary files from Figshare. After unzipping the files, the report can be found in the reports folder with the corresponding R markdown file.

#### Missing data visualizations

The Biosamples data frame within the HTML report focusing on putative human samples was used for data visualization of key metadata column types. For age, the 3000+ metadata columns were filtered using dplyr select(matches(“\\bage\\b”)) to filter age-related columns. For gender, the metadata columns were filtered using dplyr select(matches(“gender|sex”, ignore.case = T)) to filter age-related columns. For ethnicity, the metadata columns were filtered using dplyr select(matches(“ethnicity|race”)) to filter ethnicity-related columns. For body mass index (BMI), the metadata columns were filtered using dplyr select(matches(“\\bheight\\b|\\bweight\\b|\\bbmi\\b”, ignore.case = T)) to filter BMI-related columns. Metadata-filtered data frames were then used in gg_miss_var function call to visualize the missing data of each type.

## Conflict of Interest Statement

The Bioinformatics and Computational Biosciences Branch developed the Metagenote and Nephele tools described in this article. All views in this manuscript reflect the Authors’ personal views and do not necessarily reflect those of the National Institutes of Health.

## Funding

This project has been funded in part with Federal funds from the National Institute of Allergy and Infectious Diseases (NIAID), National Institutes of Health, Department of Health and Human Services under BCBB Support Services Contract HHSN316201300006W/HHSN27200002 to MSC, Inc.

## Acknowledgements

We would like to acknowledge Dr. Lynn Schriml, Dr. Robb Finn, and Dr. Lorna Richardson for helpful discussions and feedback. This research was supported in part by the Office of Science Management and Operations (OSMO) of the NIAID.

## Notes

https://doi.org/10.6084/m9.figshare.13918190.v1

